# Identification of *in vivo* CFTR conformations during biogenesis and upon misfolding by covalent protein painting (CPP)

**DOI:** 10.1101/2021.03.02.433670

**Authors:** Sandra Pankow, Casimir Bamberger, Salvador Martínez-Bartolomé, Sung-Kyu Park, John R. Yates

**Affiliations:** Department of Chemical Physiology, The Scripps Research Institute, 10550 North Torrey Pines Road, La Jolla, CA 92037, USA

**Keywords:** Chemical footprinting, high resolution mass spectrometry, misfolding, conformational defects

## Abstract

*In vivo* characterization of protein structures or protein structural changes after perturbation is a major challenge. Therefore, experiments to characterize protein structures are typically performed *in vitro* and with highly purified proteins or protein complexes. Using a novel low-resolution method named Covalent Protein Painting (CPP) that allows the characterization of protein conformations *in vivo*, we are the first to report how an ion channel, the Cystic Fibrosis Transmembrane Conductance Regulator (CFTR), is conformationally changed during biogenesis and channel opening in the cell. Our study led to the identification of a novel opening mechanism for CFTR by revealing that the interaction of the intracellular loop 2 (ICL2) with the nucleotide binding domain 2 (NDB2) of CFTR is needed for channel gating, and this interaction occurs concomitantly with changes to the narrow part of the pore and the walker A lysine in NBD1. However, the ICL2:NBD2 interface, which forms a “ball-in-a-socket” motif, is uncoupled during biogenesis, likely to prevent inadvertent channel activation during transport. In particular, solvent accessibility of lysine 273 (K273) in ICL2 changes with the opening and closing of the channel. Mutation of K273 severely impaired CFTR biogenesis and led to accumulation of CFTR in the Golgi and TGN. CPP further revealed that, even upon treatment with current approved drugs or at permissive temperature, the uncoupled state of ICL2 is a prominent feature of the misfolded CFTR mutants ΔF508 and N1303K that cause Cystic Fibrosis (CF), which suggests that stabilization of this interface could produce a more efficient CF drug. CPP is able to characterize a protein in its native environment and measure the effect of complex PTMs and protein interactions on protein structure, making it broadly applicable and invaluable for the development of new therapies.

## Introduction

The tertiary structure of a protein is closely linked to its function and determines how it interacts with other proteins. Mutations generating conformational defects are frequently associated with loss- or gain-of-function of a protein, and result in a variety of protein misfolding diseases such as CF or Alzheimer’s ^1-3^. Additionally, many of the proteins that cause misfolding diseases contain either long stretches of inherently disordered regions, are membrane proteins, or both, presenting a significant challenge to methods that depend on the determination of high-resolution structures to understand the conformational defects.

Recent cryo-EM studies of the human Cystic Fibrosis Transmembrane Regulator (CFTR), an anion channel important for maintaining the salt balance in stratified epitheliums such as the lung and intestine, have provided insight into channel architecture and structural rearrangements occurring during gating. However, the structures of several CFTR mutants that cause Cystic Fibrosis, the most common genetic childhood disease in Caucasians, have still not been solved, mainly due to technical problems to express and purify enough of these unstable proteins. In fact, even the wt CFTR structure could only be obtained by introducing stabilizing mutations and deleting a small disordered region, the RI element, which is physiologically important for CFTR maturation and is highly phosphorylated *in vivo* ^4^. To overcome these constraints, we have developed covalent protein painting (CPP), a novel structural proteomics approach that allows identification and quantification of protein misfolding events *in vivo*.

CPP covalently labels the epsilon amine of a lysine side chain *in vivo*. Lysine side chains contain primary amines that are positively charged at physiological pH. They occur primarily on exposed surfaces of the native tertiary structure of a protein where they are readily accessible for interaction with other proteins as well as for labeling with amine-reactive reagents ^5^. If lysines are involved in a protein interaction, they may be masked for labeling as they become part of the protein binding interface and therefore become solvent-excluded ^6^. Thus, labeling reagents can be used to map protein structure and interactions by measuring the differences in the availability of the amino acid side chain to react with a label. These techniques have been successfully employed in top down approaches ^5^ for peptide mass mapping to identify modification sites and provide site-specific structural information on purified proteins or simple purified protein complexes.

## Results

To characterize CFTR conformations *in vivo*, surface exposed lysines in human bronchial epithelial cells expressing wt CFTR (HBE16o-) were covalently labeled with isotope encoded dimethyl groups directly in-dish for 10 min on ice (CPP) ^7^ (Fig 1A). Subsequently, cells were lysed and CFTR was co-immunoprecipitated using the CoPIT protocol ^8^. Labeled CFTR and its interactors were then digested into peptides with lysine-insensitive chymotrypsin. Subsequently, we used a second isotope combination to label the lysines that were solvent excluded in the cell but accessible after digestion of the proteins into peptides. Using quantitative mass spectrometry, labeled lysines present in a complex cell lysate can be directly identified and quantified and their solvent accessibility determined. Twenty-six lysine sites in CFTR were quantified in 3 different biological replicates and mapped onto the CFTR Cryo-EM structure (PDB 5UAK) ^9^ showing a good general agreement between solvent accessibility determined by Cryo-EM and solvent accessibility determined by CPP (Fig. 1B,1C; Table 1). For example, lysines located on the outside of the CFTR molecule such as Lys 283 were all more than 98 % solvent accessible. In addition, CPP was able to quantify the solvent accessibility of lysines in intrinsically unstructured regions such as the Regulatory Region (R-region), which is not resolved in Cryo-EM structures and the Regulatory Insertion (RI) element, which has to be deleted in recombinant CFTR used for Cryo-EM structural studies for reasons of stabilization and yield ^9^.

**Table 1.** Mapping and comparison of lysine solvent accessibilities to Cryo-EM structures. Quantified lysines and respective peptide sequences were mapped onto the available Cryo-EM structures 5UAK and 5UAB and solvent accessible isosurfaces probed with a rolling probe.

| Position | sequence | Mean surface accessibility (%) |  | Wt Cryo-EM structure | Wt cryo-EM structure, ATP-bound, phosphorylated |
| --- | --- | --- | --- | --- | --- |
| | | wt | $\Delta F508$ | | |
| K52 | QIPSVDSADNLSEKL | nd | 92.6 | accessible (93.75 Å<sup>2</sup>) | accessible (13.39 Å<sup>2</sup>) |
| K95 | LGEVTKAVQPL | 96.1 | 77.0 | partially accessible (3.51 Å<sup>2</sup>) | accessible (35.78 Å<sup>2</sup>) |
| K273 | VITSEMIENIQSVKAY | 61.8 | 95.0 | inaccessible (0.48 Å<sup>2</sup>) | partially accessible (6.92 Å<sup>2</sup>) |
| K283 | CWEEAMEKMIENLRQTEL | 98.7 | 98.9 | accessible (53.98 Å<sup>2</sup>) | accessible (39.15 Å<sup>2</sup>) |
| K370/K377 | DSLGAINKIQDFLQKQEY | 99.8 | 98.9 | accessible (41.45 Å<sup>2</sup>, 39.11 Å<sup>2</sup>) | accessible (29.92 Å<sup>2</sup>, 36.11 Å<sup>2</sup>) |
| K411 | not present in CryoEM |  |  |  |  |
| K420 | not present in CryoEM |  |  |  |  |
| K442 | SLLGTPVLKDINF | 99.2 | 95.7 | accessible (39.53 Å<sup>2</sup>) | accessible (47.84 Å<sup>2</sup>) |
| K464 | AVAGSTGAGKTSLL | 99.7 | 93.0 | accessible (9.59 Å<sup>2</sup>) | partially accessible (4.36 Å<sup>2</sup>) |
| K536 | AEKDNIVLGEGGITL | 98.8 | 94.7 | accessible (51.94 Å<sup>2</sup>) | accessible (26.99 Å<sup>2</sup>) |
| K584 | GYLDVLTEKEIF | nd | 91.9 | accessible (8.91 Å<sup>2</sup>) | accessible (18.03 Å<sup>2</sup>) |
| K1060 | ESEGRSPIFTHLVTSLKGLW | 99.0 | 98.5 | accessible (49.05 Å<sup>2</sup>) | partially accessible (3.15 Å<sup>2</sup>) |
| K1080 | FETLFHKAL | 99.9 | 99.7 | accessible (59.33 Å<sup>2</sup>) | accessible (35.59 Å<sup>2</sup>) |
| K1165 | KFIDMPTEGKPT | 99.5 | 96.6 | accessible (19.66 Å<sup>2</sup>) | accessible (54.57 Å<sup>2</sup>) |
| K1218 | TAKYTEGGNAILENISF | nd | 89.3 | accessible (41.39 Å<sup>2</sup>) | accessible (30.56 Å<sup>2</sup>) |
| K1292 | GVIPQKVF | nd | 94.4 | accessible (48.20 Å<sup>2</sup>) | partially accessible (3.01 Å<sup>2</sup>) |
| K1302 | RKNLDPYEQWSDQEIW | 98.4 | 97.9 | accessible (26.64 Å<sup>2</sup>) | accessible (13.86 Å<sup>2</sup>) |
| K1317 | KVADEVGLRSVIEQFPGKLDF | 99.7 | 99.5 | accessible (37.29 Å<sup>2</sup>) | accessible (53.09 Å<sup>2</sup>) |
| K1334 | RSVIEQFPGKLDF | 99.9 | nd | accessible (56.92 Å<sup>2</sup>) | accessible (49.60 Å<sup>2</sup>) |
| K1420 | LVIEENKVRQY | 99.3 | 94.8 | accessible (51.71 Å<sup>2</sup>) | accessible (51.65 Å<sup>2</sup>) |
| K1429 | DSIQKLL | 99.9 | 99.5 | accessible (57.27 Å<sup>2</sup>) | accessible (33.64 Å<sup>2</sup>) |

**Figure 1.**
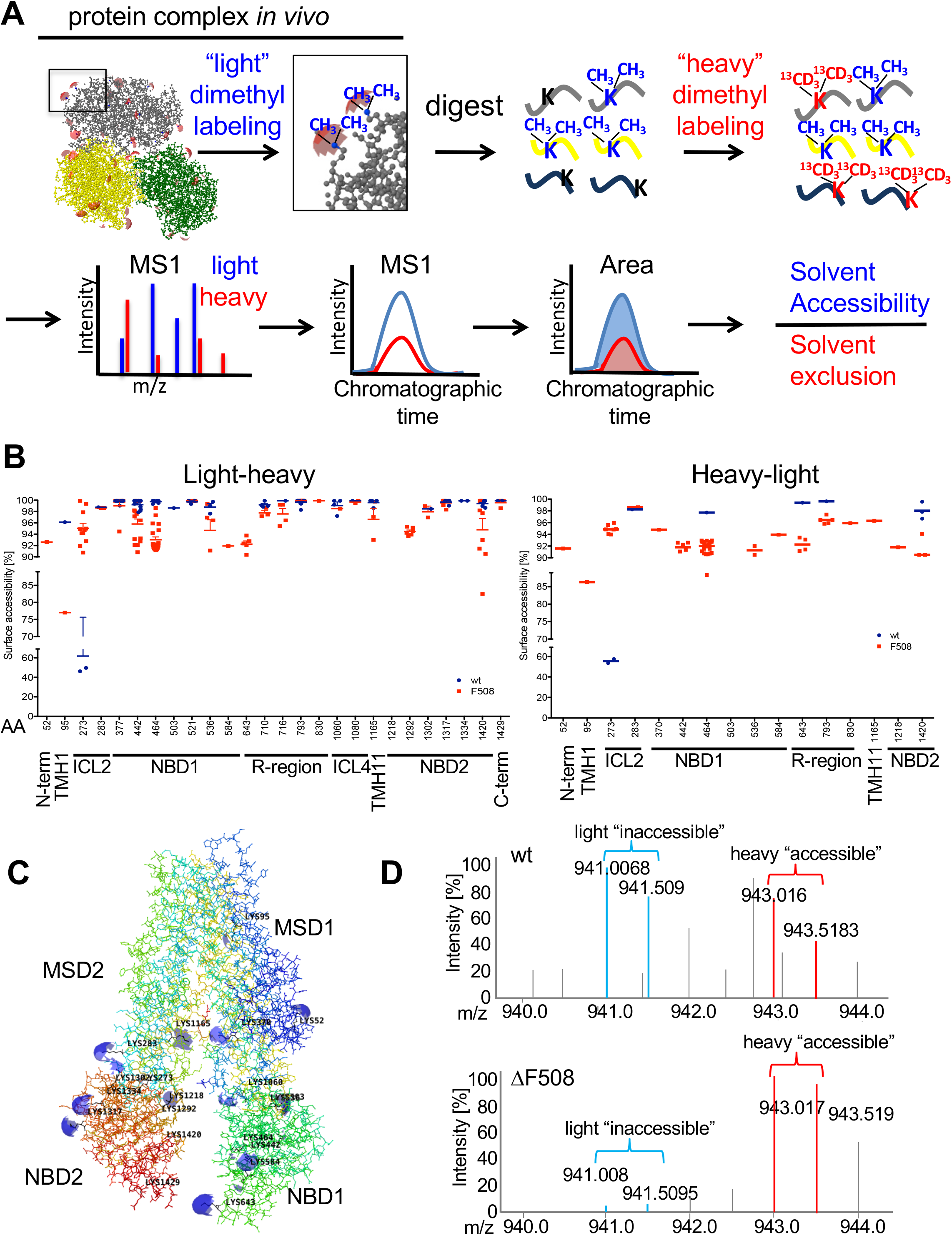
CPP analysis of wt and ΔF508 CFTR. **A**. Experimental and mass spectrometric workflow. **B**. Solvent accessibility of quantified lysines in different CFTR domains upon labeling with CH_3_(“light”) followed by ^13^CD_3_(“heavy”) labeling (left panel) and upon label-swap (right panel). **C**. Quantified lysines were mapped onto the CFTR Cryo-EM structure (PDB:5UAK) and solvent accessible isosurfaces displayed as blue spheres. **D**. Representative MS1 spectra of K273 depicting light and heavy precursors used for quantification in wt CFTR (upper panel) or ΔF508 CFTR (lower panel). Wt CFTR n=4; ΔF508 CFTR n=4 (biological replicates).

We then performed CPP on isogenic CFBE41o-cells expressing ΔF508 CFTR and compared its solvent accessibilities with wt CFTR. Our experiment revealed large differences in solvent accessibility in the transmembrane helix 1 (TMH1) and the intracellular loop 2 (ICL2) as well as smaller, but significant differences in the RI element, the N-terminus of the nucleotide binding domain 1 (NBD1) and the NBD2 (Fig.1B, left panel). A label swap experiment confirmed the detected differences between wt and ΔF508 CFTR (Fig. 1B, right panel). Particularly interesting changes were observed for Lys 95 (K95) and Lys 464 (K464). Lys 95 (K95) is part of the narrow region of the internal vestibule and is fully accessible in wt CFTR and K464 is the Walker A lysine in NBD1 and involved in ATP binding; both were less accessible in ΔF508 CFTR. However, the largest difference was observed for K273 in the ICL2, which was solvent-excluded in 40% of wt CFTR molecules, but solvent accessible to greater than 95% in ΔF508 CFTR (Fig. 1D). The ICL2 is a short “coupling helix” that reaches into the NBD2 in what has been called a “ball-in-a-socket” motif ^10^. K273 is located at the bottom of the loop and is solvent excluded in all published Cryo-EM structures as well as in 40 % of wt CFTR by CPP. Solvent accessibility of K273 thus represents a CFTR conformation in which ICL2 does not reach into NBD2, e.g. an uncoupled state. Interestingly, 60 % of wt CFTR is also in such an uncoupled conformation.

To gain a better understanding of the observed differences between wt and ΔF508 CFTR, we performed CPP experiments on ΔF508 CFTR after treatment with 5 µM of the FDA approved VX-809 CF drug^11^. We also performed CPP experiments on ΔF508 CFTR at permissive temperature of 28°C, a temperature at which ΔF508 CFTR is efficiently trafficked to the plasma membrane and partially active ^12^ (Figure 2). Treatment with VX-809 did not significantly change solvent accessibility of lysines in ΔF508 CFTR except for lysine 1218, which is located in NBD2 and is ubiquitinated in ΔF508 CFTR ^4,13^. However, permissive temperature of 28°C reduced solvent accessibility of Lys273 in ICL2 to 89% (e.g. by 6 %), showing that coupling of ICL2 to NBD2 now occurred in about 11% of CFTR molecules. We therefore speculated that ICL2 coupling to NBD2 may be crucial for CFTR activity. Comparing solvent accessibility of K273 in Cryo-EM models of inactive CFTR ^9^(PDB 5UAK) and activated, dimerized CFTR ^14^ (PDB 5UAR) revealed that K273 was solvent excluded in both models as well as in additional homology models ^15^.

**Figure 2.**
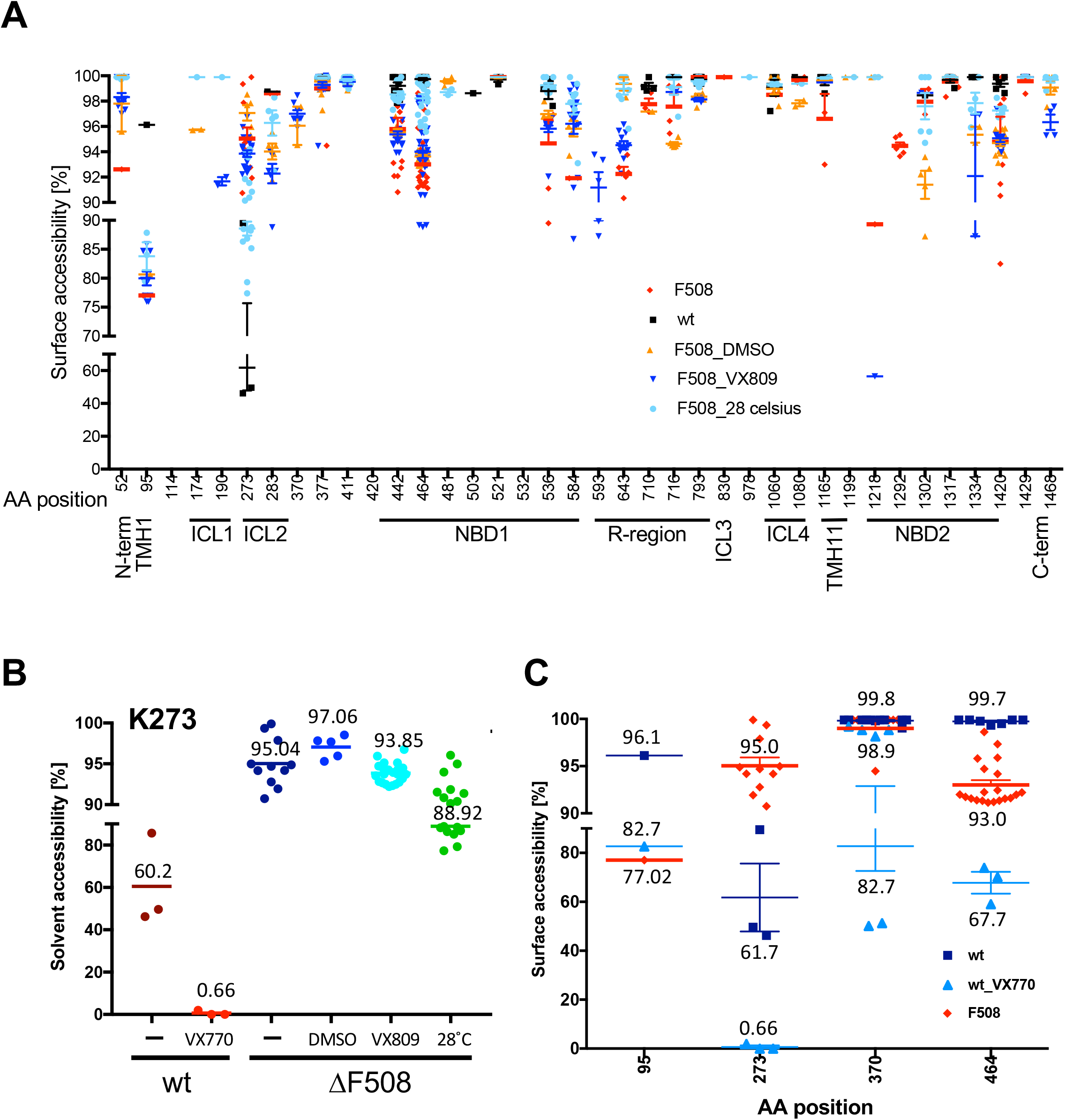
Solvent accessibility of CFTR lysines in ICL-2 and the ion permeation pathway upon correction and activation. **A**. Changes in solvent accessibility at permissive temperature of 28 °C (24h), upon VX-809 treatment (5 µM, 36h) or in DMSO control. **B**. Channel activation by VX-770 (500 nM, 10 min) changes solvent accessibility of K273 in ICL-2. Mean is indicated for each condition. **C**. Changes in solvent accessibility of lysines located in the ion permeation pathway and ATP binding site upon channel activation with VX-770. Wt (n=4), wt_VX770 (n=3), ΔF508 (n=4), ΔF508_VX809 (n=3), ΔF508_DMSO (n=3), ΔF508_28°C (n=3). n represents number of biological replicates.

To investigate the possibility that *in vivo* activation may be different from *in vitro* activation of CFTR, we treated wt CFTR with 500 nM VX-770 for 10 min^16^ and then measured K273 accessibility by CPP. VX-770 is a channel potentiator that opens CFTR channels independent of ATP hydrolysis and is used to treat CF caused by the G551D mutation ^17-19^. Surprisingly, activation of wt CFTR by VX-770 treatment led to complete solvent exclusion of K273. Only in 0.66 % of CFTR molecules was K273 still solvent accessible, thus showing a clear correlation of K273 accessibility with wt CFTR activity (Fig. 2B). Coupling of ICL2 to NBD2 upon activation with VX-770 was accompanied by additional surface accessibility changes in the vestibule (K95), the chloride pathway (Lys 370) and the Walker A lysine Lys 464, all of which became less accessible (Fig. 2C, Supplemental Figure 1A). The data suggested that the CFTR conformation in which K273 is solvent excluded and the ICL2 reaches into NBD2 is an active open state, while the conformation in which K273 is solvent accessible represents an inactive state in which ICL2 is uncoupled from NBD2.

To probe whether this previously undescribed inactive CFTR conformation is associated with gating of the channel or is rather an inactive confirmation attained by CFTR when it is not present at the plasma membrane, we performed additional CPP experiments with CFTR variants N1303K CFTR and well as G551D CFTR. N1303K CFTR, like ΔF508 CFTR, is misfolded and not present at the plasma membrane, and G551D CFTR is trafficked normally to the plasma membrane, but has a gating defect ^20-23^. CPP results on these CFTR variants showed that solvent availability of K273 was high (>95%) in the mainly ER and Golgi located, misfolded CFTR variants N1303K and ΔF508 CFTR, but that it was solvent excluded in G551D, suggesting that this conformation is not associated with CFTR gating per se (Figure 3A,3B). To investigate when this CFTR conformation occurs, we mutated CFTR K273 to K273A (which is a small, polar, cavity creating amino acid) and analyzed biogenesis of K273A CFTR by confocal microscopy. Confocal images showed that the K273A CFTR mutant exits the ER, but does not reach the plasma membrane; instead it accumulates in the cells (Figure 4A). However, these are not aggregates, as shown by missing colocalization with autophagy markers such as SQSTM and cleaved LC-3 (Supplemental Figure 2). Interactome analysis of K273A CFTR versus a GFP control followed by GO enrichment analysis revealed that proteins localized to membrane bounded organelles and vesicles were highly enriched as interactors of K273A CFTR as well as proteins involved in endosomal transport and vesicle mediated transport (Fig. 4B, 4C, Supplemental table 1). Examples of these interactors include the ESCRT complex, and the retromer cargo-selective complex (CSC), specifically the SNX3-retromer. The ESCRT complex is required for endosomal sorting of protein cargo into multivesicular bodies (MVBs)^24,25^, and the SNX3-retromer mediates retrograde transport from the endosome to the trans-Golgi-network (TGN), and transport from endosome to plasma membrane; it was also previously shown to promote cell surface expression of ENAC sodium channels ^26-28^ (Fig. 4D). However, interactors that recognize misfolded proteins in the ER and belong to the ER associated degradation (ERAD) pathway and are over-represented in ΔF508 CFTR interactomes, were not enriched. CPP analysis of the K273A CFTR mutant further showed that solvent accessibility of lysines was similar to wt CFTR in non-activated state (Figure 5D).

**Figure 3.**
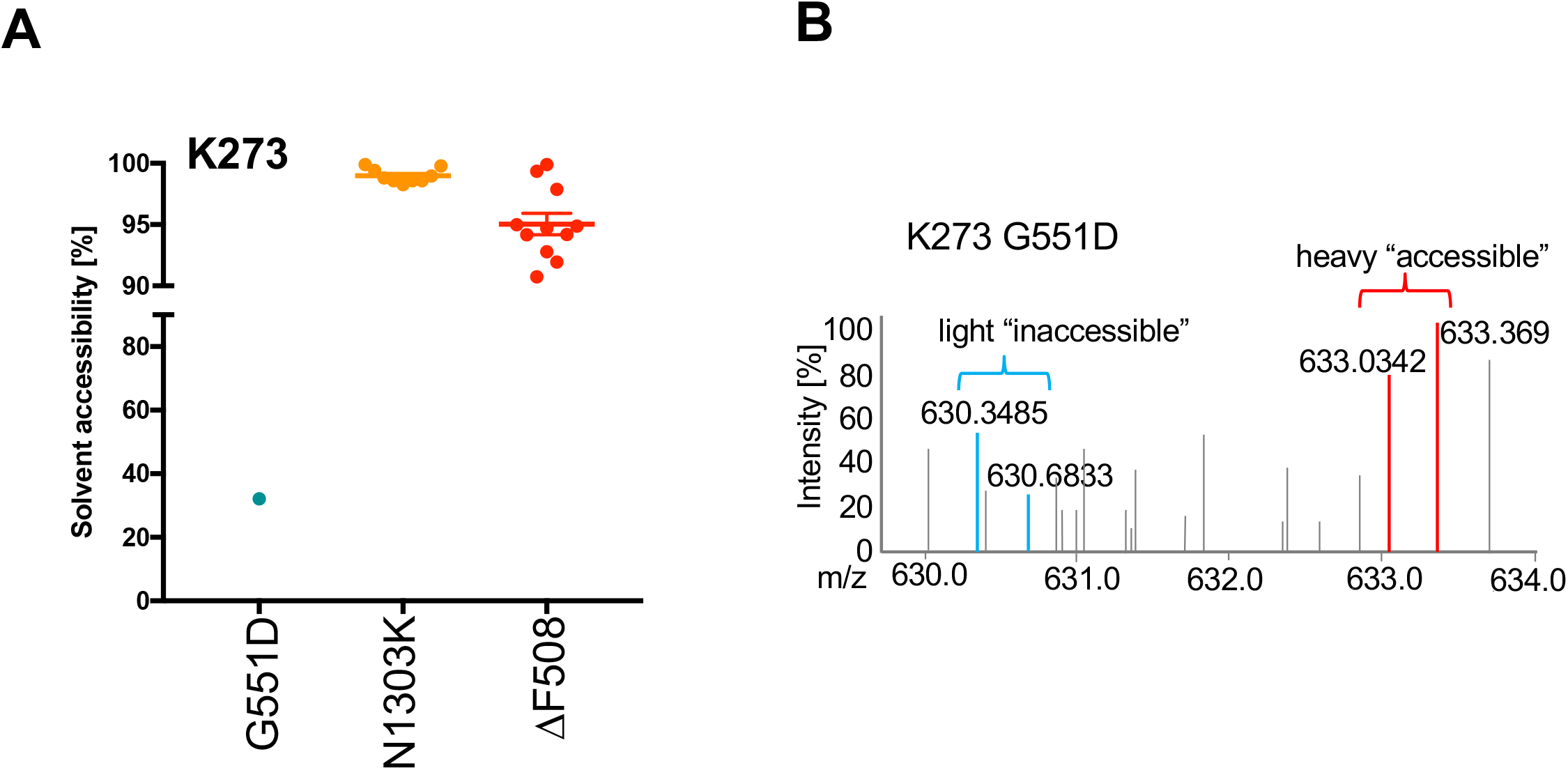
K273 solvent accessibility in G551D and N1303K CFTR. **A**. Solvent accessibility of K273. **B**. Representative MS1 spectrum of the peptide containing K273 in G551D CFTR. Light and heavy precursors and measured m/z for each are indicated. G551D (n=3), N1303K (n=3). n represents biological replicates.

**Figure 4.**
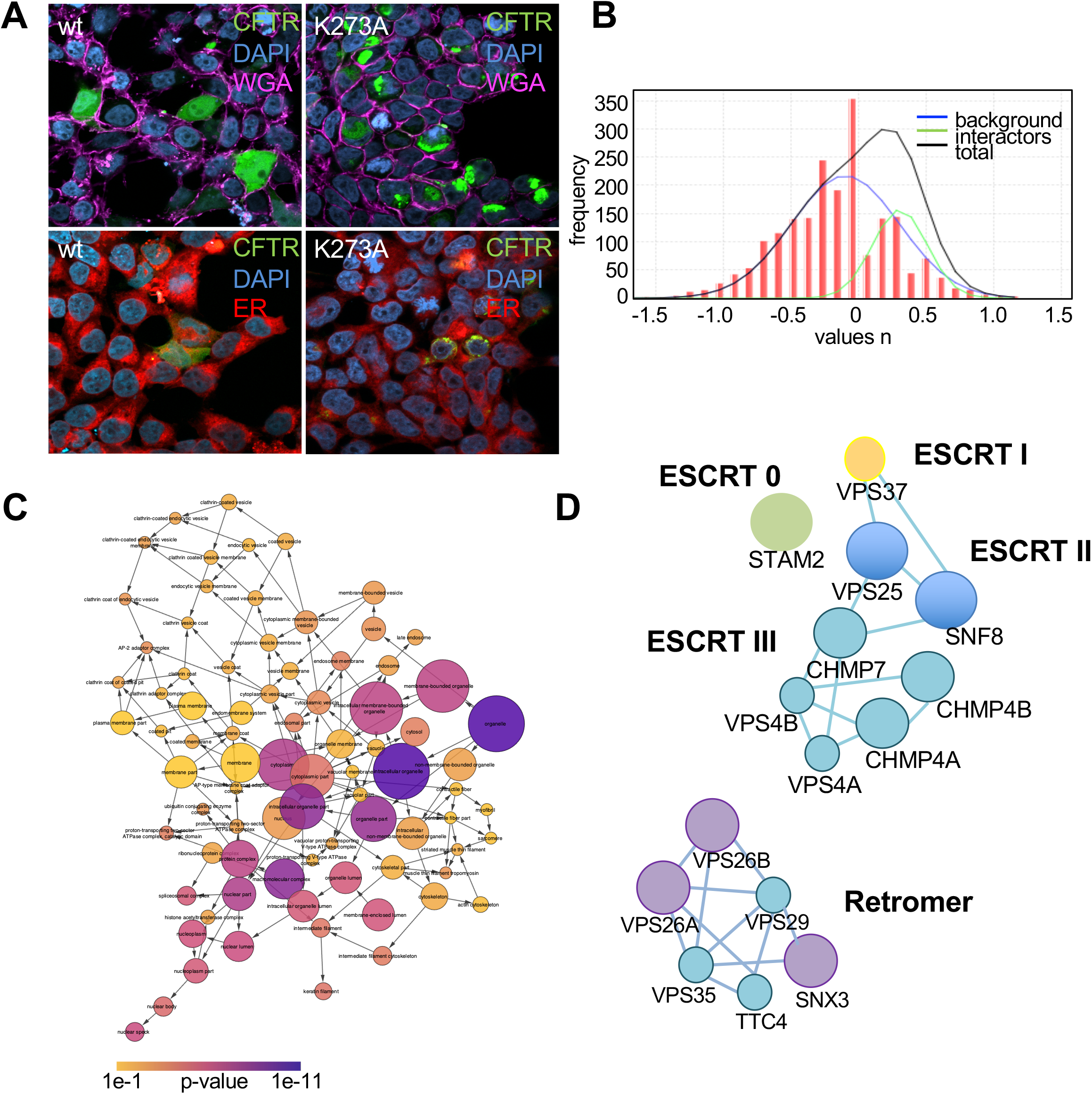
K273A CFTR interactome and cellular localization. **A**. HEK293T cells transiently transfected with GFP tagged wt or K273A CFTR (green) were stained with wheat germ agglutinin (pink) to visualize the plasma membrane (upper panel). ER was stained with the ER Cytopainter kit (lower panel). Nuclei were counterstained with DAPI (blue).n=3 biological replicates. **B**. Interactome analysis of K273A CFTR. True K273A CFTR interactors were distinguished from background using CoPITgenerator. **C**. Network of enriched GO-terms in the K273A CFTR interactome. **D**. ESCRT and retromer complexes were specifically enriched in the K273A interactome.

## Discussion

Here, we report a novel CFTR conformation that reflects an immature, closed state of the channel. This conformation is attained by 60 % of wt CFTR molecules in the cell as well as the misfolding mutants ΔF508 CFTR and N1303K_[JY1][MOU2]_. The data also suggest previously unknown conformational rearrangements that appear upon activation of the channel in the ion permeation pathway and which add important detail to the knowledge gained from Cryo-EM structures.

The most striking feature of the newly discovered immature CFTR conformation is the altered solvent accessibility of K273 in the ICL2 coupling helix. In general, ABC transporter ICLs mediate the coupling of ATPase (catalytic) activity at the nucleotide binding sites to substrate translocation through the membrane spanning domains (MSDs) ^29^. Previous data suggested that ICL2 is involved in stabilization of the full conductance state of the CFTR Cl channel, as a 19 amino acid deletion mutant of CFTR ICL2 is non-responsive to forskolin stimulation and does not reach the cell surface ^30^. In models of the ABC transporter P-glycoprotein the ICLs may move like a ball in a socket to accommodate conformational change during the gating cycle ^10^. In this arrangement, the ICL2 loop reaches into the NBD2 as in all current Cryo-EM structures of CFTR and thus is not solvent accessible. Such solvent exclusion of K273 was observed upon activation of the CFTR channel with VX-770. Interestingly, crosslinking cysteines engineered in CFTR’s ICL2 and NBD2 decreased the open probability of the channel (*P*_*o*_) dramatically ^31,32^, suggesting that movement of the ICL2 is needed for proper channel gating. However, if K273 is fully solvent accessible as it is in ΔF508, N1303K and 60% of wt and G551D CFTR, it must represent a conformation in which the ICL2 is removed from the socket in NBD2 (i.e., an “uncoupled” state), and thus represents an inactive CFTR conformation.

One could interpret the different conformations simply as open/closed states during the gating cycle. However, the fact that this inactive conformation is also observed in ΔF508 and N1303K CFTR, two CFTR variants that are misfolded and are largely degraded before reaching the plasma membrane, suggests otherwise. Furthermore, based on its interactome and confocal images, K273A mutation led to accumulation of CFTR in the Trans-Golgi network (TGN), ERGIC and endosomes. This indicates that the inactive conformation is an immature CFTR conformation that is attained during transport from the ER to the plasma membrane and may prevent accidental channel activation during biogenesis and transport. Thus, CPP was used to discover an immature CFTR conformation that was previously unobservable. It is possible that this conformation is stabilized by interactors that prevent ICL2 from reaching into NBD2 or by internal molecular re-arrangements, such as movement of the R-domain or the RI element upon phosphorylation. While movement of the R-domain from its place between the two NBDs to the peripheral surface has been observed in the PKA phosphorylated, ATP-bound CFTR ^14^, it cannot be the only factor that prevents the coupling of ICL2 to NBD2, as ICL2 also reaches into NBD2 in the structure of the un-phosphorylated, ATP-free CFTR. We therefore suspect that additional *in vivo* phosphorylation events are needed, or that movement of the RI element (which is deleted in all CFTR structures) is involved since it is highly phosphorylated in wt CFTR, but not in ΔF508 CFTR and the amount of RI element phosphorylation correlates with the amount of fully mature CFTR in the plasma membrane ^4^.

The CPP results also present an opportunity to further examine changes in the ion permeation pathway that occur upon channel activation. Changes include the walker A lysine K464 becoming less solvent accessible upon VX-770 activation, likely reflecting ATP-binding and dimerization of the NBDs. This is consistent with the observation that NBD binding symmetrically closes off the ATP-binding sites in the activated state ^14^. Further changes include K95 and K370, two charged residues in the vestibule and close to the chloride entry pore ^9^, which become partially inaccessible in activated wt CFTR treated with VX-770 and the G551D mutant. This could reflect a percentage of channels that have closed, or may reflect conformational changes that reflect regional pore constriction/tightening at the mouth of the pore and the vestibule. Regional pore constriction/tightening is consistent with a role of these positively charged residues in anion selection and in creating an electrostatic potential for chloride ions to move along the pore ^33^. Comparing CPP results across all analyzed CFTR mutants and upon activation, we also noticed changes in solvent accessibility of K52, which is part of the “lasso motif” and a structure unique to CFTR ^34^. K52 was accessible in ΔF508 and N1303K, but only partially accessible in G551D and in VX-770 activated wt CFTR, suggesting a role of the lasso in channel activation or conformational changes occurring upon localization to the plasma membrane. Such a role would be consistent with previous observations that the lasso motif is important for interactions with the membrane traffic machinery and would confirm suggestions that the lasso regulates CFTR channel gating through interactions with the R domain ^14,35-37^.

Finally, the CPP data revealed a few changes that appear to be unique to N1303K, ΔF508 and G551D CFTR and thus related to the misfolding and the gating defect, respectively. In comparison to ΔF508 CFTR, N1303K CFTR exhibited higher solvent accessibility at K584 in NBD1 and K643 located at the border of the NBD1 and R-region. The G551D mutant exhibited a specific difference in solvent accessibility of K536. A change unique to ΔF508 CFTR was the slightly altered solvent accessibility of K1420 in NBD2. Interestingly, structural differences between wt and ΔF508 CFTR in NBD2 have been previously identified by limited trypsin digestion assays ^38^. Finally, CPP data also allowed us to better rationalize a VX-809 corrector mechanism, which remains largely unknown. VX-809 has been reported to bind to NBD1 using isolated NBD1 and to stabilize domain:domain interfaces, particularly NBD1:ICL4, although others observed that VX-809 alone do not stabilize isolated NBD1 ^39-41^. In this study, we observed a reduction in solvent accessibility of K1218 by more than 30%, a very clear change that was not observed in any other condition (Supplemental table 2). Point mutation of K1218 to K1218R has been shown to increase the amount of CFTR, stabilize it at the plasma membrane and reduce lysosomal degradation of CFTR ^13^, while its deletion improves folding efficiency of ΔF508 CFTR ^42^. Binding of VX-809 to the region or K1218 itself could thus diminish ΔF508 CFTR targeting to the lysosome, although it is unlikely to completely prevent it, as multiple mechanism exist for peripheral quality control of CFTR ^43,44^. Such a mechanism would be in line with the moderate rescue of ΔF508 CFTR processing observed for VX-809.

## Methods

### Cell culture, plasmids and point mutations

CFBE and HBE cells were grown in A-MEM supplemented with 10% FBS, 1% Penicillin-Streptomycin (GibCo, Carlsbad, CA) and appropriate antibiotics at 37°C, 5% CO_2_. FRT cells expressing either wt CFTR or the variants G551D, N1303K or ΔF508 CFTR were cultured in Ham’s F12 medium, Coon’s modification supplemented with 5 % FBS and 100 µg/ml Hygromycin. K273A CFTR was generated using the Quick change method (Quiagen) in the pCMV6-ac-CFTR-GFP plasmid (Origene). Hek293T cells were transfected with Lipofectamine 3 according to manufacturer’s recommendation (Thermo Fisher, Carlsbad, CA) and expression of K273A CFTR was analyzed 48 h to 72 h after transfection.

### In vivo dimethyl labeling and CFTR-IP

Cell culture media was removed and the intact cell layer was washed with 1x PBS before labeling of surface-exposed lysines with either formaldehyde (CH3, 0.3 % final concentration) and sodiumcyanoborohydrate (NaBH3CN, final concentration 30 mM) for light labeling or with deuterated formaldehyde containing ^13^C (^13^CD3) and deuterated sodiumcyanoborohydrate (NaBD_3_CN) for heavy labeling. The labeling reaction was allowed to continue for 10 min on ice before it was stopped by adding 1% ammonium acetate (NH 4C_2_H_3_O). Subsequently, cells were harvested with a cell scraper and lysed by adding 2 x TNI buffer ^8^ containing complete Ultra protease inhibitor (Roche) and Halt phosphatase inhibitor (Pierce). The cell lysate was sonicated for 3 min in a water bath sonicator and insoluble material was removed by centrifugation at 18,000 x g, 4°C, 15 min. CFTR-IP was then performed as described in Pankow et al., 2016 ^8^, except that the last three washes were performed with HNN buffer (50 mM Hepes, pH 7.5, 150 mM NaCl, 1 mM EDTA). Proteins were precipitated by methanol-chloroform precipitation (Lysate: Methanol: Chloroform, 1:4:1, *v*:*v*:*v*) and the precipitate was washed with 95% Methanol before digestion. For interactome analyses, CFTR was immunoprecipitated as described ^45^ using GFP-Trap Agarose (Chromotek).

### Protein digestion and peptide labeling

Precipitated proteins were resolubilized by sonication in 0.2% Rapigest, 100 mM HEPES, pH 8.0 in a water bath sonicator. The protein lysate was heated to 95°C for 10 min, reduced by 5 mM TCEP (Pierce), and alkylated with 10 mM iodoacetamide (Sigma-Aldrich, St.Louis, MO) before digestion with chymotrypsin (Protea Biosciences, Morgantown, WV, 1:100 w:w) at 30°C for 14-16 hours. Peptides were labeled according to Boersema et al., 2009 ^46^ with 0.16% CH3 or ^13^CD3 and 30 mM NaBH3CN or NaBD_3_CN, respectively, for 1 h at room temperature. The labeling reaction was stopped with 1% NH_4_C_2_H_3_O, Rapigest was inactivated by incubation with 9% formic acid (1h, 37°C) and samples were reduced to near dryness *in vacuo*. Tryptic digestion of unlabeled CFTR IPs was carried out as described previously ^8^.

### Mass spectrometry

Peptides were reconstituted in buffer A (95% H_2_O, 5% acetonitrile/0.1% formic acid), loaded onto a preparative MudPIT column and analyzed by nano-ESI-LC/LC-MS/MS on an LTQ-Elite or LTQ-VelosPro Orbitrap (Thermo Fisher, San Jose, CA) by placing the triphasic MudPIT column in-line with an Agilent 1200 quaternary HPLC pump (Agilent, Palo Alto, CA) and separating the peptides in multiple dimensions with a 10-step gradient (0%, 10%, 20%, 30%, 40%, 50%, 60%, 70 %, 80%, 90% Buffer C ((500 mM ammonium acetate/5% acetonitrile/0.1% formic acid)) over 20 h as described previously ^47^. To avoid cross contaminations between different samples, each sample was loaded onto a fresh column. Each full scan mass spectrum (400-2000 m/z) was acquired at 60,000 R, followed by 20 data-dependent MS/MS scans at 35% normalized collisional energy and an ion count threshold of 1000. Dynamic exclusion was used with an exclusion list of 500, repeat time of 60 s and asymmetric exclusion window of -0.51 and +1.5 Da. Monoisotopic peaks were extracted from Raw-files with RawConverter ^48^ and MS/MS spectra were searched with ProLuCID ^49^ against the human Uniprot database (release Nov 2019) using a target-decoy approach in which each protein sequence is reversed and concatenated to the normal database ^50^. The search was carried out using a search window of 50 ppm, no enzyme specificity, N-terminal static modification according to protein dimethyl label (e.g. 28.0313 or 36.0757), and lysine dimethylation (28.0313) and carbamidomethylation of cysteine (57.02146) as static modifications. Additionally, oxidation of methionine (15.9949) and serine, threonine or tyrosine phosphorylation (79.9663) were considered as differential modifications. A lysine mass shift of 8.0444 was used for metabolic labeling search. Minimum peptide length was set to 7 amino acids (AA). Search results were filtered to less than 0.2 % FDR on spectrum level, corresponding to less than 0.5 % peptide level FDR using DTASelect 2.1 within the Integrated Proteomics Pipeline (IPA, San Diego, CA). Additionally, a deltaCN value of ≥ 0.1, and precursor delta mass cutoff of less than 8 ppm were applied as filter.

### Quantification of solvent-excluded vs solvent-exposed lysines and mapping of solvent accessible areas

To determine the proportion of solvent-exclusion to solvent-exposure of a lysine residue, the area under each peptide peak was quantified using Census and the Integrated Proteomics Pipeline (IPA, San Diego, CA) as described previously ^51,52^. Individual isotopes were extracted within a 10 ppm window. A determinant factor of ≥ 0.5, a profile score of ≥ 0.5 and no retention time shift allowed were set as parameters to filter for correct peak assignments. The determined ratio of light over heavy directly reflects solvent-exposure/ solvent-exclusion. Peak shapes were also manually assessed for large ratio measurements. Protein structures available in PDB were visualized with JMOL (version 14.30.1) and the isosurface (isoSurface saSurface) of the epsilon amino group of lysine calculated and visualized in JMOL with a rolling probe radius of 1.2 Å. Area ratios were expressed as percentage of surface accessibility as described ^7^ and plotted in Prism 7 (Graphpad Inc.) Statistical analysis of area ratios per site was carried out in PCQ as described ^53^. Briefly, ratio values for each lysine site were calculated with the SoPaX algorithm, which is part of ProteinClusterQuant (PCQ, https://github.com/proteomicsyates/ProteinClusterQuant), a One-way ANOVA analysis carried out and obtained results subjected to Benjamini-Hochberg correction.

### Immunofluorescence staining and western blotting

Cells were grown in Lab Tek II chambers (Thermo Fisher), fixed with 4 % paraformaldehyde/1xPBS for 10 min at RT and solubilized with 0.1% Triton X-100 /1xPBS before staining with antibodies against LC3A/B (Cell Signaling Technologies, cat no. 12741S) and SQSTM1 (Cell Signaling Technologies, cat no. 8025S). The plasma membrane was visualized by wheat germ agglutin CF 594 conjugate (Biotium) and the ER was visualized using the ER Cytopainter kit (Abcam) according to manufacturer’s recommendations. Slides were mounted in ProLong Gold Antifade Mountant with DAPI to visualize nuclei (Thermo Fisher). Images were taken with a Zeiss LSM 710 confocal laser scanning microscope. Protein lysates were prepared in TNI buffer as described previously^8^. CFTR was detected using mouse monoclonal antibodies 24.1 (ATCC) and M3A7 (EMD Millipore). Antibodies against b-actin (AC-15, Sigma) and Na+/K+ATPase a (H300, sc28800, Santa Cruz) were used for normalization. Horseradish-peroxidase-conjugated secondary antibodies (Jackson Immunoresearch) were detected with enhanced chemiluminescence (ECL, Pierce).

## Author contributions

S.P., C.B. and J.R.Y. designed the research and S.P. and C.B. performed the experiments. S.M.B. implemented the protein residue-specific quantification and SoPaX in PCQ. S.K.P. supported mass spectrometric data analysis and developed quantification software. J.R.Y. provided materials and funding. S.P. wrote the manuscript and prepared the figures with help from all authors. All authors read and approved the manuscript.

## Acknowledgements

We would like to thank E. Sorscher (University of Alabama) for the kind gift of the FRT cell lines and Claire Delahunty (TSRI) for reading the manuscript and editing it for clarity. Funding was provided by NIH grants R01HL131697-05 and R33CA212973-01 awarded to John R. Yates III.

## Figure legends

**Supplemental Figure 1.**
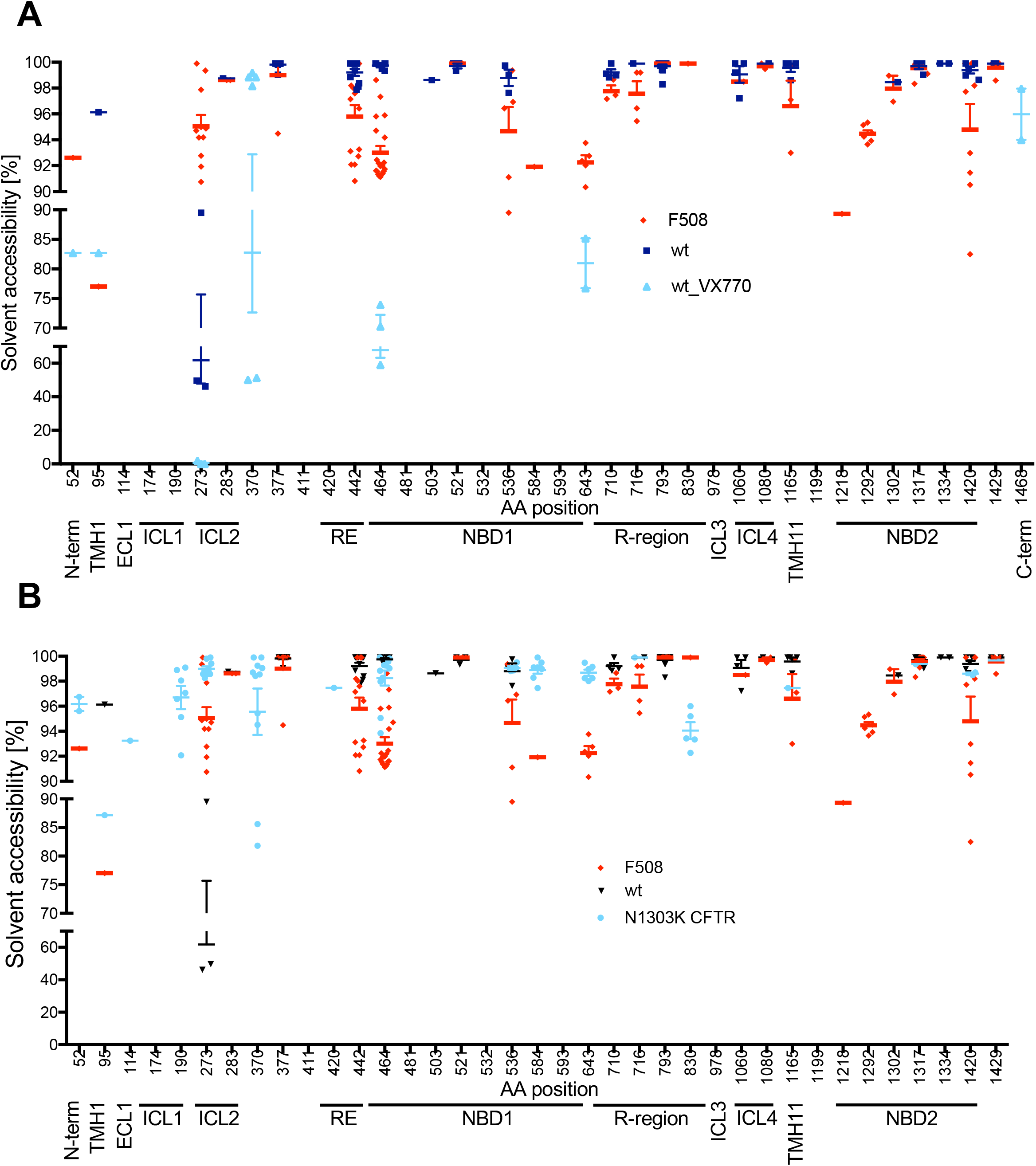
**A**. Solvent accessibility of all quantified CFTR lysines upon activation with VX-770 (n=3). **B**. Solvent accessibility of all quantified N1303K CFTR lysines compared to ΔF508 and wt CFTR. N1303K (n=3).

**Supplemental Figure 2.**
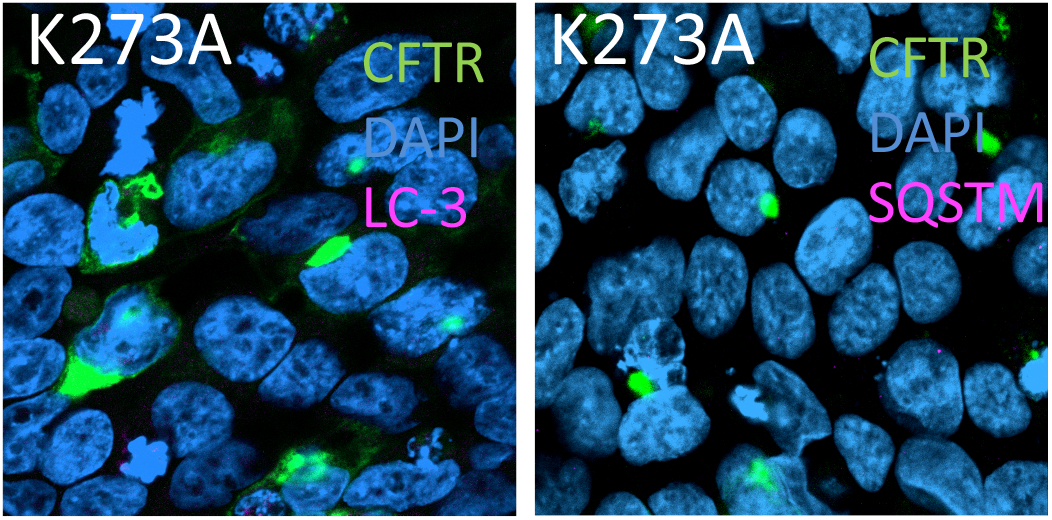
HEK293T cells transiently transfected with GFP-tagged wt or K273A CFTR were stained for autophagy markers SQSTM (pink) or cleaved LC-3 (pink).

**Supplemental table 1**. Enriched GO-terms in K273A interactome with p-values and adjusted p-values.

**Supplemental table 2**. Means and standard deviation (SD) of all quantified lysines, the identified peptide sequences and CFTR AA position are displayed for all measured conditions.

